# A Dual-Functional Needle-Based VOC Sensing Platform for Rapid Vegetable Quality Examination

**DOI:** 10.1101/2024.12.12.628229

**Authors:** Oindrila Hossain, Yan Wang, Mingzhuo Li, Sina Jamalzadegan, Noor Mohammad, Alireza Velayati, Aditi Dey Poonam, Qingshan Wei

## Abstract

Volatile organic compounds (VOCs) are common constituents of fruits, vegetables, and crops, and are closely associated with their quality attributes, such as firmness, sugar level, ripeness, translucency, and pungency levels. While VOCs are vital for assessing vegetable quality, traditional detection methods, such as Gas Chromatography-Mass Spectrometry (GC-MS) and Proton Transfer Reaction Mass Spectrometry (PTR-MS) are limited by expensive equipment, complex sample preparation, and slow turnaround time. Additionally, the transient nature of VOCs complicates their detection using these methods. Here, we developed a paper-based colorimetric sensor array combined with needles that could induce vegetable VOC release in a minimally invasive fashion and analyze VOCs *in situ* with a smartphone reader device. The colorimetric sensor array was optimized using sulfur compounds as main targets and classified fourteen different vegetable VOCs, including sulfoxides, sulfides, mercaptans, thiophenes, and aldehydes. By combining principal components analysis (PCA) analysis, the integrated sensor platform proficiently discriminated between four vegetable subtypes originating from two major categories within 2 min of testing time. This rapid and minimally invasive sensing technology holds great promise for conducting field-based vegetable quality monitoring.

**Graphical abstract:** 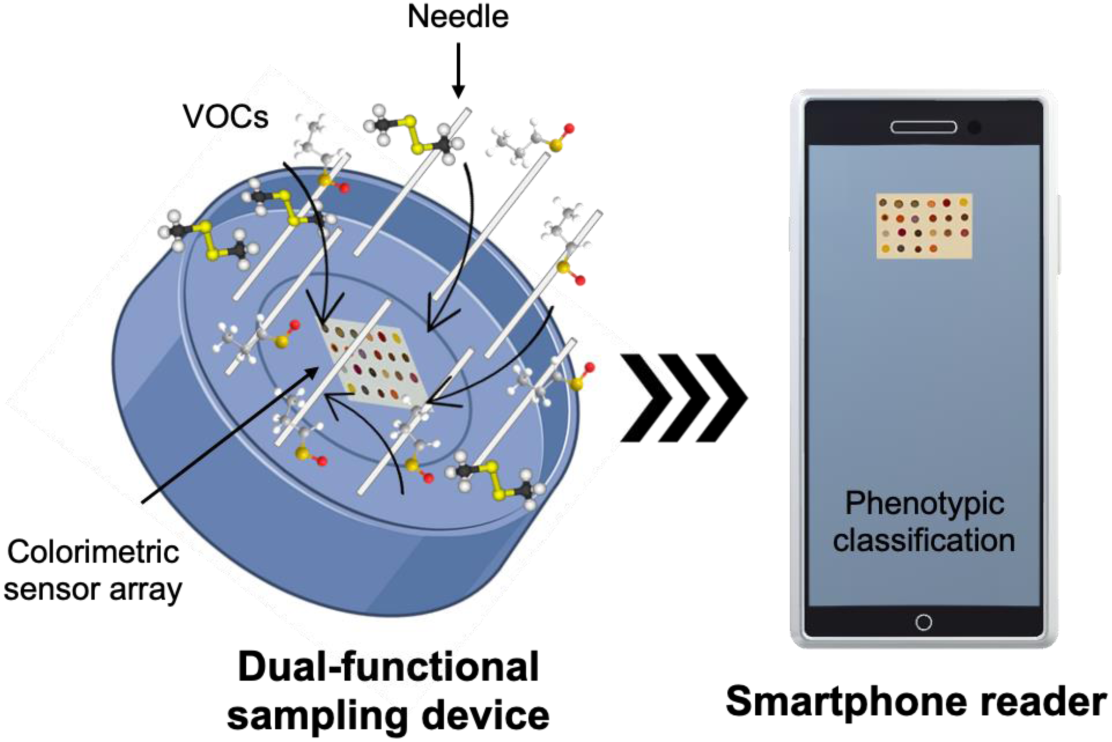

## 1. Introduction

Volatile organic compounds (VOCs) serve as essential indicators in both realms of plant disease detection and monitoring of vegetable and fruit quality ( Gao et al. 2022; Li et al. 2021; Li et al. 2019; Mei et al. 2023; Zhao et al. 2022). In the context of plant health, these organic compounds act as messengers of distress, which are emitted in a unique pattern when plants are infected by pathogens or under environmental stresses (Jones et al. 2016; Wu et al. 2023). This makes VOC analysis a powerful tool for noninvasive and early disease detection, enabling farmers to take proactive measures to protect their crops (Lee et al. 2023; Li et al. 2019; Tholl et al. 2021). Furthermore, VOCs are equally instrumental in the assessment of vegetable and fruit quality (Janssens et al.; Lu et al. 2012; Qiu and Wang 2015). The presence and concentration of specific VOCs can signify ripeness, flavor, and overall freshness, making them invaluable in quality control processes across the agricultural industry (Gao et al. 2022; Passa et al. 2023; Taiti et al. 2015; Tiwari et al. 2020; Wang et al. 2021). Phenotypic classification of vegetables and fruits is an integral part of the quality monitoring process. Knowing the subtypes of a particular fruit or vegetable is essential for variety monitoring during the breeding process. The distinct chemical profiles of different vegetable and fruit species can be identified through VOC analysis, contributing to accurate phenotypic classification (Gao et al. 2022).

The VOCs present in vegetables or fruits constitute a complex mixture of chemical compounds that contribute to the wide range of aromas and flavors in these edible plants (Gao et al. 2022; Taiti et al. 2015; Tiwari et al. 2020). Common VOCs in vegetables/fruits include alcohols, aldehydes (responsible for grassy or nutty notes), and sulfur-containing compounds that give some vegetables characteristic pungency (Cecchi et al. 2020; Gao et al. 2022; Passa et al. 2023; Taiti et al. 2015; Tiwari et al. 2020; Zhang et al. 2017). Esters can also be found, adding fruity or floral hints to certain vegetables (Cuevas et al. 2017; Mei et al. 2023). Previous studies suggest that a few common VOCs detected in vegetables are propanethiol, dimethyl disulfide, acetone, hexanal, Ethyl butanoate, 1-hexanol, and 1-octen-3-ol (Gao et al. 2022; Mathieu et al. 2009; Wang et al. 2021; Zhang et al. 2022; Zhang et al. 2021a). However, vegetable VOCs may decompose quickly after releasing or during the sample preparation steps, creating major challenges for accurate VOC characterization with lab-based instrumentation methods. Since the volatile profile from vegetables is highly dependent on the timing of sampling, the conventional liquid phase extraction method and gas chromatography-mass spectroscopy (GC-MS) are not suitable for rapid VOC capture and time-resolved quantification. Instead, solid phase microextraction (SPME) has been utilized to effectively trap volatiles, which can then be quantified by direct insertion into the gas chromatography (GC) instrument (Gao et al. 2022). Moreover, real time-mass spectrometry (DART-MS) and proton-transfer reaction-mass spectrometry (PTR-MS) have been demonstrated to detect the volatiles for sensitive, selective, and fast measurement (Lindinger and Jordan 1998; Zhang et al. 2021b). Nevertheless, all these methods require extensive time, labor, and sophisticated laboratory equipment to perform the analysis. In addition, another major disadvantage of these methods is their invasive sampling nature, which renders vegetables unsuitable for consumer distribution after testing.

The advancements in portable VOC sensors have revolutionized the ways to monitor food quality, air quality, and plant health ( Endo et al. 2007; Lee et al. 2023; Li et al. 2019; Lim et al. 2009; Taoukis and Tsironi 2016; Tholl et al. 2021; Xiao-wei et al. 2016). These sensors have significantly improved the ability to detect and quantify VOCs in real-time or on-site without much human operation. Previously, we developed a field-portable noninvasive VOC detection system based on a smartphone to detect late blight in tomatoes (Li et al. 2019). To the best of our knowledge, there currently exists no handheld VOC sensing platform capable of fruit/vegetable quality monitoring. It is challenging because many vegetables exhibit very weak VOC emissions while intact. Some vegetables do not produce certain specific VOCs until mechanical induction such as cutting is performed. Thus, there arises an imperative need for the development of a device capable of both VOC induction and detection, facilitating on-site detection without losing instantaneous VOCs.

Here, we present a novel VOC sensing system that utilizes a needle-integrated colorimetric sensor device to differentiate phenotypic classes of various vegetable varieties, while minimizing the damage to the targets. The colorimetric sensing array consists of a wide range of chemically responsive colorants, such as metalloporphyrins and different chromatic dyes belonging to various families such as metal salts, pH indicators, solvatochromic/vapochromic dyes, and metal ions. With a sensitivity level in the ppm range, the colorimetric sensor distinguished various sulfur-containing VOCs associated with vegetables. The needles on the device minimally invaded the vegetables and induced the VOC release. The colorimetric sensor on the same device responded to the emitted VOCs by changing the color, which was captured by a smartphone-based reader. The sensor system effectively differentiated and categorized four varieties of vegetables with high classification accuracy when tested in the lab environment.

## 2. Materials and methodology

### 2.1 Reagents and materials

All reagents and materials were analytical reagent grade and used without further purification. Reagents for colorimetric sensor array preparation (**Table S1**) were purchased from Millipore Sigma. The cellulose ester membranes (catalog no. HF18004XSS) were purchased from EMD Millipore. A thermoplastic ABSplus-P430 (Eden Prairie) was used to make smartphone attachments and sensor cartridges by 3D printing.

### 2.2 Preparation of the image-capturing device

An Android smartphone (LG V10) was used to capture the colorimetric strip images. The smartphone attachment design was similar to our previously developed design with slight modifications (removing the pump) (Li et al. 2019).

### 2.3 Sensing material preparation for liquid phase experiments

To prepare the organically modified sol-gel (ormosil) formulation, tetramethyl orthosilicate (TMOS), methyltrimethoxysilane (MTMS), methanol, and milli-Q water were mixed in the molar ratio of 1:1:11:5. The mixture was stirred at room temperature for 2 h to complete the ormosil preparation (Lim et al. 2008). The sensing material’s ID numbers are followed in **Table S2**.

Sensing materials 1-3: Pyrrolidine amine (2 x 10^-2^ M) was added to a solution of Zn-tetraphenylporphyrin (2 x 10^-3^ M) in Chloroform (CHCl_3_). The amine and porphyrin solution concentrations were modified from the original protocol to enhance the color change (Leontiev and Rudkevich 2005). The same steps were followed for the preparation of Tetrakis(2,4,6-trimethylphenyl)porphyrinatocobalt(II)-pyrrolidine and Tetrakis(pentafluorophenyl)porphine iron(III)chloride-pyrrolidine sensing materials.

Sensing materials 4-7: Various amounts (1-5 mg) of methyl red were added to 500 μL of sol-gel to adjust the solubility. The pH of the methyl red solution was adjusted by adding different amounts of base (0.1-8 μL 1M Sodium hydroxide). Similar adjustments were made for nitrazine yellow, phenol red, and cresol red solution preparation.

Sensing material 8: Pararosaniline hydrochloride was dissolved in a 10% aqueous ethanol solution to make a pararosaniline stock solution (3.088 × 10^−3^ mol/L). The stock solution, 36% formaldehyde solution, and 32% hydrochloride acid solution were mixed in a molar ratio of 1:125:1255, respectively (Monro et al. 2012). The final solution was completed by adding 2.2905 mL water. The amount of water was modified to make a visible spot so that it could be seen on the paper substrate.

Sensing material 9: Two different amines, such as Methyl diethanolamine (MDEA) and pyrrolidine, were mixed separately with cresol red. 2 x 10^-3^ M cresol red solution was prepared by adding 0.81 mg dye in 1 mL CHCl_3_. 1.67 μL MDEA was added to the dye solution. Similarly, pyrrolidine was added to a separate dye solution. The final composition of the sensing material was modified from the original protocol (Chatterjee and Sen 2015).

Sensing material 10-12: m-cresol purple dye was mixed with 500 μL of sol-gel by adjusting the solubility of the dye in the sol-gel solution. The amount of TBAH base solution (1M Tetrabutylammonium hydroxide in 2-methoxyethanol) was also adjusted (1-5 μL). Two different salts (silver nitrate and zinc acetate) were chosen to mix with the basic dye solution.

Sensing material 13-14, 16-22: These dyes mentioned in **Table S2** were added individually to the sol-gel solution. The amount of dye was determined based on the solubility and visibility by the naked eye.

Sensing material 15: 4-(4-nitrobenzyl) pyridine was mixed with N-benzylaniline in a 1:1 mass ratio and added to 500 μL sol-gel solution. 10 μL of each sensing material solution was transferred to the coverslip (Fisher Scientific Catalog No.50-143-780), and images of the droplet were taken. After that, 1 μL of target analyte was added to it. Another image was captured. Similar steps were followed for several targets, as mentioned in **Table S2**.

### 2.4 Sensor array preparation

Supplementary Table S3. contains comprehensive information regarding the sensing materials and concentration of individual sensor components. A slotted stainless-steel pin (parts no. FP4CB, V&P Scientific) was dipped in each sensing element formulation and printed on the cellulose ester membrane. Each sensing material formed a round shape spot ∼1 mm in diameter. Then the strips were oven-dried at 60℃ overnight and further dried for another 24 h at 35℃. The colorimetric strips were stored in a glass petri plate within a nitrogen-filled desiccator. The Petri plate was wrapped with parafilm (Bemis) and aluminum foil to reduce air and light interference, respectively.

### 2.5 Vapor phase experiment and sensor reproducibility test

Sensors were incubated with each organic target analyte (total of fifteen analytes) for 1 h in a glass petri dish and wrapped with parafilm to generate saturated vapor from the liquid analyte. The sensor images before and after exposure were captured by an LG V10 smartphone camera. To determine the optimum exposure time, sensor strips were incubated with 1-pentanethiol, diethyl sulfide, and dimethyl sulfoxide for 5 min, 15 min, 30 min, 45 min, and 60 min. The strip images were taken at each time point of the measurement. For the limit of detection experiment, a gas generation unit was used to prepare different concentrations of volatiles (5-1000 ppm) (Cellini et al. 2017). Briefly, two mass flow controllers (MKS instruments GM50A013204SBM020 and Alicat™ Scientific MC-500SCCM) were used to dilute the saturated vapor of the analyte by mixing nitrogen gas at a specific ratio. The gas mixture flowed over the colorimetric sensors kept in a glass bowl, maintaining the continuous flow of the gas. Similarly, the photos were taken by a smartphone after 30 min of exposure. For the control experiments, nitrogen flowed over the strips without the volatiles.

### 2.6 Vegetable VOCs detection experiments

After cutting and grinding the vegetable, 1.25 g puree was weighed in a small petri plate. Then, the sample was incubated with three sensor strips in a bigger petri plate and wrapped with parafilm (**Figure 4a**). The incubation of the vegetable sample and colorimetric sensors was continued from 15 min to 16 h. All images of the colorimetric strips were taken by a smartphone.

### 2.7 Needle device preparation

Several holders were tested to prepare the needle device, such as bottle caps, centrifuge tube caps, and microneedle patches. Blunt needles were attached to the holder by drilling with a flexible rotating option. The colorimetric sensor strip was pasted on the inside of the holder with double-sided chemical-resistant tape.

### 2.8 Needle-induced VOC detection experiment

After pressing the colorimetric sensor-loaded needle device on the vegetable surface, it was left on the vegetable for a certain time (2 min, 5 min, 15 min, or 30 min) to interact VOCs with the sensor. Then, the needle device was removed from the surface of the vegetable. The smartphone took the before and after exposure images.

### 2.9 Blind human test to detect vegetable types

A blind test was conducted for ten vegetables (five Vegetable 1 and five Vegetable 2). Each vegetable was cut through the equator and two portions were held close to the face for 10-15 seconds. The results were noted based on smell and sensory irritation.

### 2.10 Data analysis and classification

For all sensing experiments, the smartphone images were used without any modifications. A MATLAB algorithm was used to detect a single spot on the sensor array and extract RGB values from the spot. The algorithm provided the difference in RGB values of the before and after-exposed images. In order to prevent the occurrence of subtraction artifacts that may arise from color variations near the edge of a spot, calculations were limited to a region of interest (ROI) (0.5 mm from the center of the spot). By averaging the ROIs of the spots, any potential artifacts resulting from non-uniform printing of the spots, particularly at the edges, were minimized. The background color of the strip (area excluding the spots) was also normalized for before and after-exposure images to minimize the color change associated with the smartphone flashlight. The color display maps were created by Microsoft Excel 365 (Microsoft Corp.). The PCA plots were generated using Python software.

## 3. Results and discussion

### 3.1 Sensor material selection and optimization

#### Sensing materials selection and screening

Previous studies showed that sulfur oxides, disulfides, thiophene, sulfides, trisulfides, thiols, aldehydes, ketones, esters, and alcohols are frequently present in vegetables (Cecchi et al. 2020; Zhang et al. 2017). As such, we designed a colorimetric sensor array comprising organic dyes mixed with additives such as amines, bases, and metal salts, targeting sulfur-based volatiles present in vegetables. **Table S1** shows the primarily selected sensing dyes. The response of the selected sensing materials towards target VOC markers was first tested using a solution-exposure experiment (**Figure 1a**), and then confirmed by the gas-exposure experiment (**Figure 1b** and **1c**).

**Figure 1.**
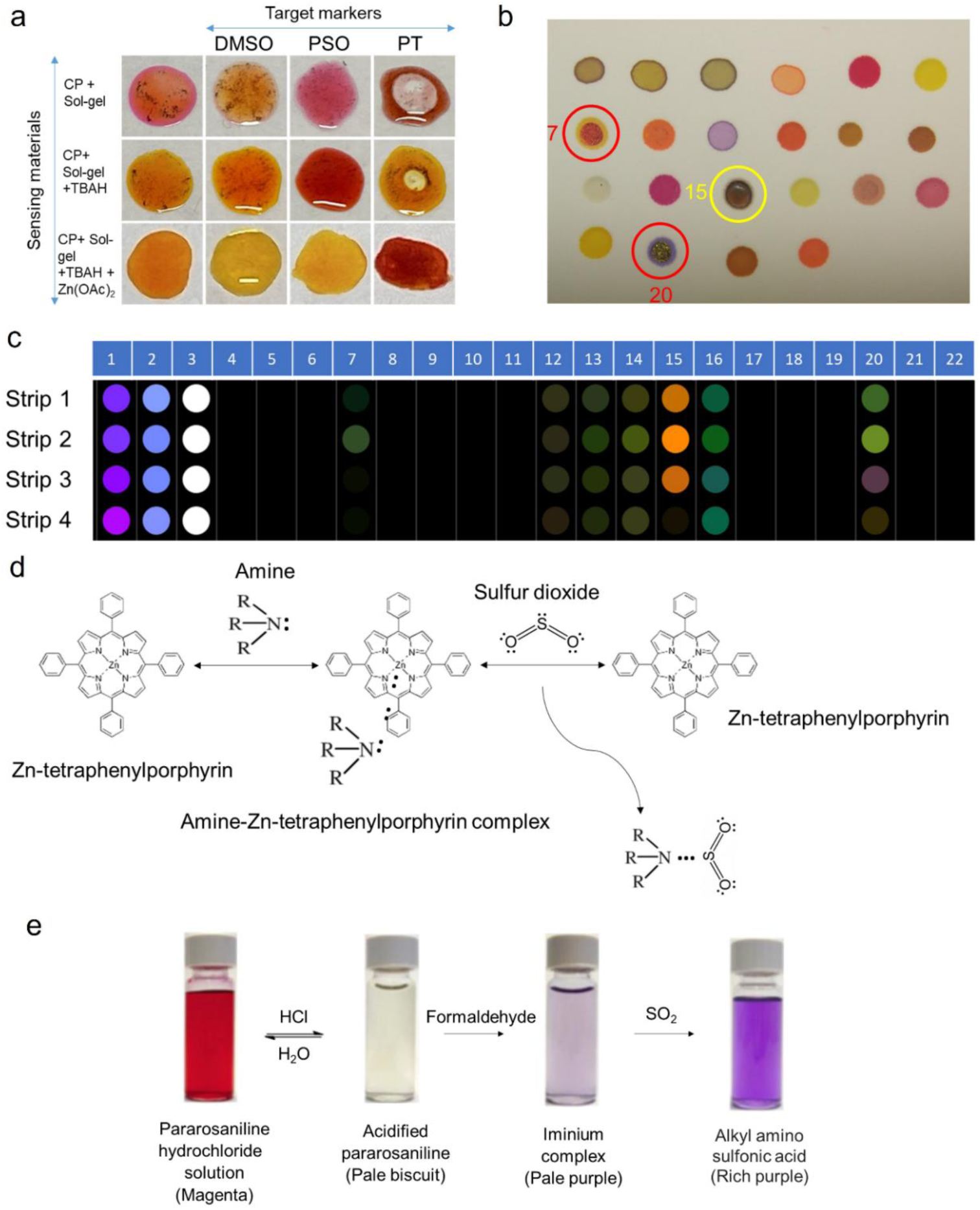
Screening of colorimetric sensing materials. (a) Sensing material solutions response to liquid target analytes; (b) A 22-element colorimetric sensor array (Spot no. 7, 15, and 20 were removed in future experiments); (c) Differential RGB colormap of 4 individual color strips exposed to 1-Pentanethiol saturated vapor (∼18,000 ppm); (d) An indicator-displacement assay for the detection of SO_2_ (Modified from Leontiev and Rudkevich (2005))(Leontiev and Rudkevich 2005); (e) Response of pararosaniline hydrochloride, HCl, and formaldehyde complex to SO_2_ (Modified from Monro *et al*. (2012)) (Monro et al. 2012). Abbreviations: CP: m-cresol purple, TBAH: Tetrabutyl ammonium hydroxide, Zn(OAc)_2_: Zinc Acetate, DMSO: Dimethyl Sulfoxide, PSO: Propyl Sulfoxide, PT: 1-Pentanethiol.

For the solution-exposure experiment, the organic dyes were prepared in a sol-gel formulation except for metalloporphyrins. The pH of the dye-containing solution was modified using bases (sodium hydroxide (NaOH) or tetrabutylammonium hydroxide (TBAH)) depending on the volatile target groups. Two different amines, pyrrolidine or methyl diethanolamine (MDEA), were used for metalloporphyrins and organic dyes, respectively, to detect sulfoxide-containing molecules.

Previous research showed that the metal salt-loaded organic dyes have a response to hydrogen sulfide (H_2_S) (Lim et al. 2009). We tested similar organic dyes and metal salts to detect thiol compounds by modifying their concentration and pH. **Figure 1a** shows the response of zinc acetate metal salt-loaded m-cresol purple (CP) organic dye to three sulfurous volatiles, namely dimethyl sulfoxide (DMSO), propyl sulfoxide (PSO), and 1-pentanethiol (PT). The dye solution composition was systematically optimized to enhance the color change. At first, m-cresol purple powder was dissolved in sol-gel, and 10 μL of this solution was pipetted on a microscope cover slip. When 1 μL of DMSO was added to the dye solution, it became pale yellow from pink. We didn’t observe any change in the dye solution after adding PSO. PT and cresol purple solution were immiscible; therefore, there was no visible color change (**Figure 1a**, top row). In the control group (**Figure 1a**, middle row), a base (TBAH) was added to the dye solution to reduce the immiscibility issue with 1-pentanethiol. In a similar way, DMSO was added to the basic dye solution, and a slight color change was observed. In contrast, a strong color change was observed for PSO: the dye solution turned from yellow to red. In the last control group, we added metal salts (silver nitrate/zinc acetate) to the cresol purple-base solution. We found that silver nitrate was not soluble in the dye solution and zinc acetate salt was a better replacement. A significant color change was visible after adding DMSO, PSO, and PT, suggesting enhanced color response. Metal salt helped PT fully dissolve in the dye solution, leading to a strong color change from yellow to deep red. The reason is that sulfur compounds reduce the metal salts into metal nanoparticles, resulting in the production of acidic by-products. These by-products can be identified using pH indicators that are integrated into the nanoporous sol-gel spot (Lim et al. 2009). **Table S2** shows the response of metal salt-containing organic dyes to several vegetable volatiles in liquid form (Sensing materials ID 10-12). From this experiment, we have determined that the metal salt is essential for the sensing materials to be responsive to thiol volatiles.

Zn-tetraphenylporphyrin-amine complex was previously used to detect sulfur dioxide (SO_2_) (Leontiev and Rudkevich 2005). First, amines react with metalloporphyrin to produce coordination complexes that exhibit color. Then, the introduction of SO_2_ induces a competition between the porphyrin and amine, which ultimately causes the release of the porphyrin and results in alterations to the color of the solution (Figure 1d) (Leontiev and Rudkevich 2005). Based on this sensing mechanism, we selected several metalloporphyrins and amines (pyrrolidine and MDEA) to detect sulfoxide compounds. The concentration of pyrrolidine and MDEA was modified from the original protocol to enhance the color intensity of the sensing material so that they are visible when printed on paper. The response of the amine-metalloporphyrins has been tested for liquid phase dimethyl and dipropyl sulfoxide, and the results are shown in **Table S2**. Because of the weakly acidic nature of dimethyl sulfoxide, it couldn’t make strong color changes in the solution, whereas dipropyl sulfoxide made substantial changes in color. Alternatively, Askim et al. used acid indicators to detect SO_2_ (Askim et al. 2016). We mixed sodium hydroxide (NaOH) with methyl red, nitrazine yellow, phenol red, and cresol red dyes in the sol-gel solution, respectively. We adjusted the base (NaOH) volume to change the pH range of the solution, which can exhibit color change for various sulfoxide-containing chemicals. The detailed composition is shown in **Table S3** (Sensing materials ID 5-8). We found that methyl red, phenol red, and cresol red were responsive to dipropyl sulfoxide but not to dimethyl sulfoxide. Monro et al. demonstrated the detection of SO_2_ in wine using a pararosaniline solution (mixture of pararosaniline hydrochloride, hydrochloric acid, and formaldehyde). The HCl-conjugated pararosaniline hydrochloride solution forms an iminium complex with formaldehyde, which changes to alkyl amino sulfonic acid in exposure to sulfur dioxide (Figure 1e) (Monro et al. 2012). The molar ratio of pararosaniline chloride solution, HCl, formaldehyde, and water was modified to detect the sulfoxide group of vegetables. It changed color from pale purple to deep purple upon adding DMSO and PSO. Finally, a previous study showed that amine-conjugated cresol red on porous aluminum support could be used for the detection of CO_2_ and SO_2_ in air (Chatterjee and Sen 2015). We used the concept of adding amine to the organic dyes to detect sulfoxide molecules. We used pyrrolidine and MDEA at several concentrations and mixed them with cresol red in sol-gel. A similar response was observed for targeted sulfoxide chemicals (DMSO, PSO) as amine-metalloporpyrins.

Solvatochromic dyes (i.e., Reichardt’s Dye, Nile red) have been previously used to detect hydrogen sulfide saturated vapor through strong dipole–dipole interactions (Suslick et al. 2004). We therefore chose several solvatochromic dyes to detect sulfide-containing volatile compounds in vegetables. The amount of Reichardt’s Dye and merocyanine 540 in sol-gel solution was adjusted based on their solubility. We found that Reichardt’s Dye was responsive to both dimethyl and diethyl sulfide. However, merocyanine 540 was responsive to only diethyl sulfide. Several mixing ratios of 4-(4-nitrobenzyl)pyridine and N-benzyniline were tested and mixed with sol-gel. This sensing material was responsive to both sulfides. 1-ethyl-4-(2-hydroxystyryl)pyridinium iodide and bromopyrogallol red changed color due to diethyl sulfide addition, but no response was observed for dimethyl sulfide. Nile red was responsive to both sulfides. The composition of the sulfide-detecting sensing materials (13-18) is described in **Table S3**.

For detecting aldehydes and ketones, several pH indicators have been used in the literature (Janzen et al. 2006). We selected thymol blue, bromophenol red, pyrocatechol violet, and cresol red, and mixed them separately with sol-gel solution to fully dissolve them, making a uniform solution of each dye. They showed an excellent response to several aldehydes such as (E)-2-hexenal, commonly mentioned as aldehydes in **Table S2**. These dyes also showed a response to dimethyl sulfoxide.

Finally, twenty-two sensing complexes were selected and printed on an ester cellulose membrane for the vapor phase validation test (Figure 1b). When the MDEA mixed metalloporphyrins and cresol red were transferred on the paper, they penetrated and spread on the paper, making a large hole due to the strong basic nature of MDEA. To reduce the problem, we replaced all the MDEA amine with relatively weak amine (pyrrolidine). Sensing materials IDs no. 7 and 20 had some shiny crystal-like particles on the spots after printing on the paper. The shininess of these dyes produced inconsistent results when exposed to several volatiles. For sensing materials ID no. 15, due to its penetration into the paper, the true color of the dye could not be measured accurately. These issues created strip-to-strip variations for spot no. 7, 15, and 20 as shown in Figure 1c. When the identical four sensor strips were exposed to 1-pentanethiol saturated vapor, spots 7, 15, and 20 showed significant differences in their response pattern from one strip to another. Therefore, we excluded these three sensing materials from subsequent experiments.

### 3.2 Sensor response to vegetable volatile biomarkers

#### Classification of model volatiles

To demonstrate the ability of the sensor array to discriminate among analytes, we chose fourteen volatile compounds to mimic vegetable VOCs. We tested one thiol (1-pentanethiol), two sulfoxides (dimethyl sulfoxide, propyl sulfoxide), two sulfides (dimethyl sulfide, diethyl sulfide), two disulfides (dipropyl disulfide, methyl propyl disulfide), one thiophene (3,4-demethyhiophene), one aldehyde ( (E)-2-hexenal), two ketones (2-methyl-3-heptanone, acetone), two esters (methyl salicylate, methyl jasmonate), and one alcohol (1-octen-3-ol) (Cecchi et al. 2020; Cuevas et al. 2017). Figure 2a shows the RGB differential colormap of the sensor array exposed to the saturated vapor of the above-mentioned analytes. The differential colormaps were generated by subtracting the pre-exposure RGB images from the post-exposure images. The sensing element ID was kept as 1-22, excluding spot no. 7,15, and 20. The 19 detection spots from each sensor array produced a 57-dimensional vector comprising RGB difference (ΔR, ΔG, ΔB) values for each analyte. The pixel intensity of the differential colormaps ranging from 3-20 was rescheduled to be the maximum and minimum for a better visual representation. A minimum intensity of 3 was applied to filter off background noise.

**Figure 2.**
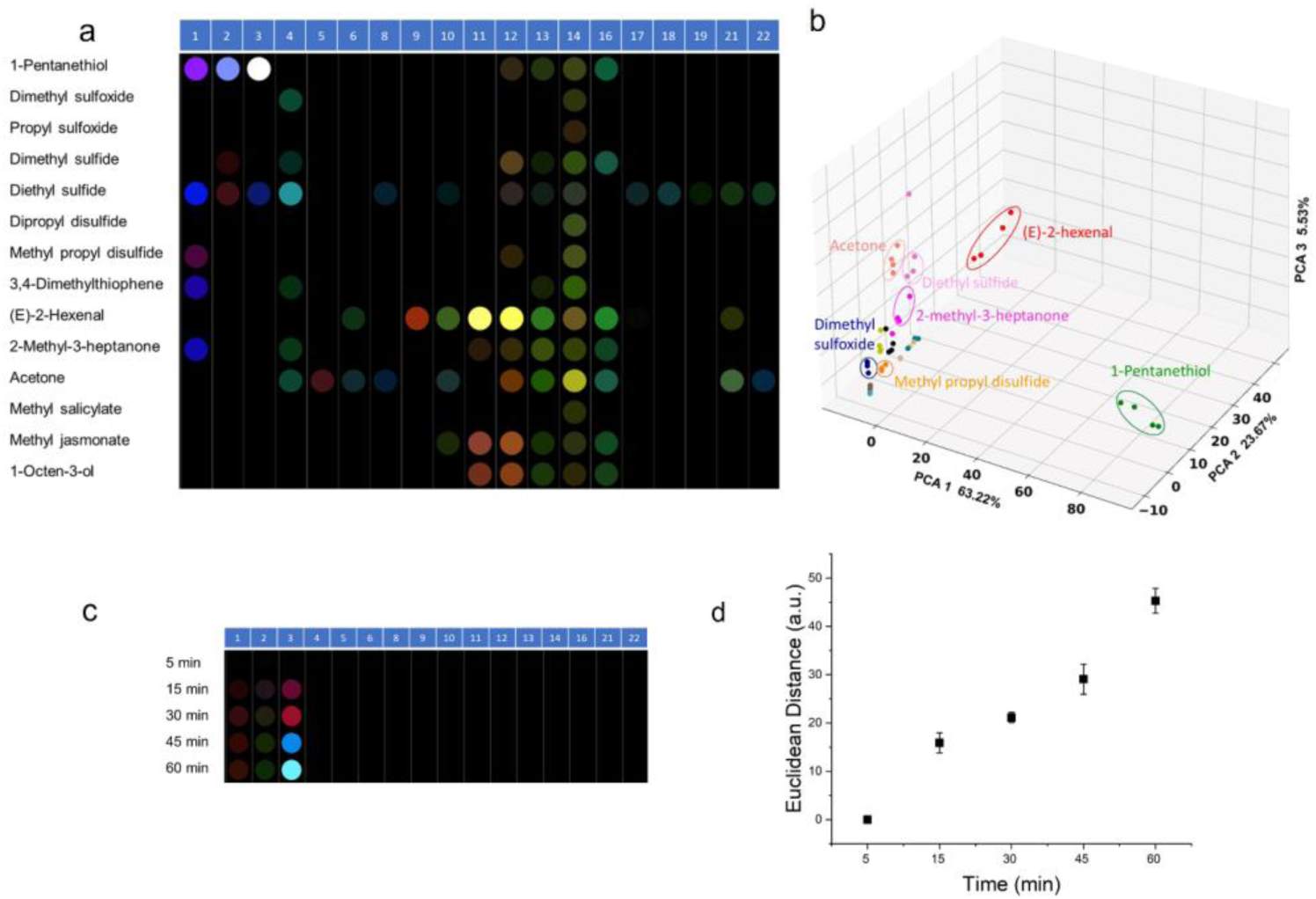
Classification of model vegetable VOCs. (a) RGB differential colormap of sensors exposed to fourteen different saturated vapor targets (each measurement is averaged over four replicates); (b) PCA plot using the first three principal components (n = 4 independent experiments); (c) RGB colormap of sensors exposed to the saturated vapor of 1-Pentanethiol at different exposure time (n = 3 independent experiments); (d) Response curve of 1-Pentanethiol vapor as a function of exposure time.

Excellent discrimination among the fourteen model volatiles was achieved visually (Figure 2a). The colorimetric sensor array was comprised of cross-reactive sensing dyes, each of which can respond to several analytes. However, the color change pattern from the whole sensor array was uniquely distinct from one target to another. We applied principal component analysis (PCA) (Kangas et al. 2018) to separate the analytes by reducing the data dimension to three for the simplicity of visualization and plotting. In the 3-axis PCA plot, ∼91% discriminatory ability was captured (Figure 2b). The well-separated clusters were observed for 1-pentanethiol, (E)-2-hexenal, acetone, diethyl sulfide, dimethyl sulfoxide, methyl propyl disulfide, 2-methyl-3-heptanone. However, alcohol (brown dots) and esters (sea green and khaki dots) were overlapped to a certain degree, which may be due to the similarity of these targets and the reduction in the data dimensionality after the PCA process (Feng et al. 2010). As sensing materials IDs no. 17, 18, and 19 were either nonresponsive or slightly responsive to the target VOCs, we excluded them from the real vegetable testing.

#### Analytical performance of colorimetric sensor array

The reproducibility, minimum exposure time, and limit of detection (LOD) of the sensor array were determined. The sensor reproducibility was tested for several analytes using four different sensor strips produced from different batches on different days. **Figure S1** shows the colormap of the four independent runs (four different sensor strips) for each analyte. The behavior of the sensor strips was similar to each other for a single analyte with a coefficient of variation in the range of ∼5-10%, showing the consistency of batch-to-batch sensor strip performance.

To investigate the minimum exposure time of the sensor to the analytes, several strong and weak analytes were incubated with sensors at different time points. Figures 2c and **Figure S2** show the RBG colormap of the 1-pentanethiol, diethyl sulfide, and dimethyl sulfoxide at 5, 15-, 30-, 45-, and 60-min exposure times. Figure 2d was plotted based on Euclidean distance (ED) (where ED=√Δ*R*^2^ + Δ*G*^2^ + Δ*B*^2^) as a function of exposure time. ED represents the absolute color shift related to the original RGB values. The results suggested that 15 min of exposure time was enough for a significant color change of the sensor for detecting a saturated vapor. A longer exposure time (30 min) could be beneficial for targets with lower vapor pressure. Therefore, to maintain the same condition for all analytes to ease the comparison, we followed 30 min exposure time for the LOD experiment.

To determine the LOD of the sensor array, a titration experiment was conducted for several model vegetable volatiles by diluting the saturated vapors with nitrogen gas. Figure 3a shows a schematic representation of the gas mixing and dilution system. Nitrogen and saturated vapor of liquid analytes were mixed at a certain flow rate to produce final target gas concentration in the range of 5-1000 ppm. Figure 3b shows the RGB color profile of 1-pentanethiol vapor at different concentrations and control (no VOC, top row) by averaging the response from three colorimetric strips. To calculate LOD, the ED values were plotted with respect to the concentrations of VOC targets in Figure 3c. The LOD was determined to be 25 ppm based on the P-value analysis. A similar experiment was conducted for 3,4-dimethylthiophene vapor (Figure 3d), and the LOD was 5 ppm (Figure 3e). All LODs are in good agreement with the reported vegetable volatile concentrations in the literature.

**Figure 3.**
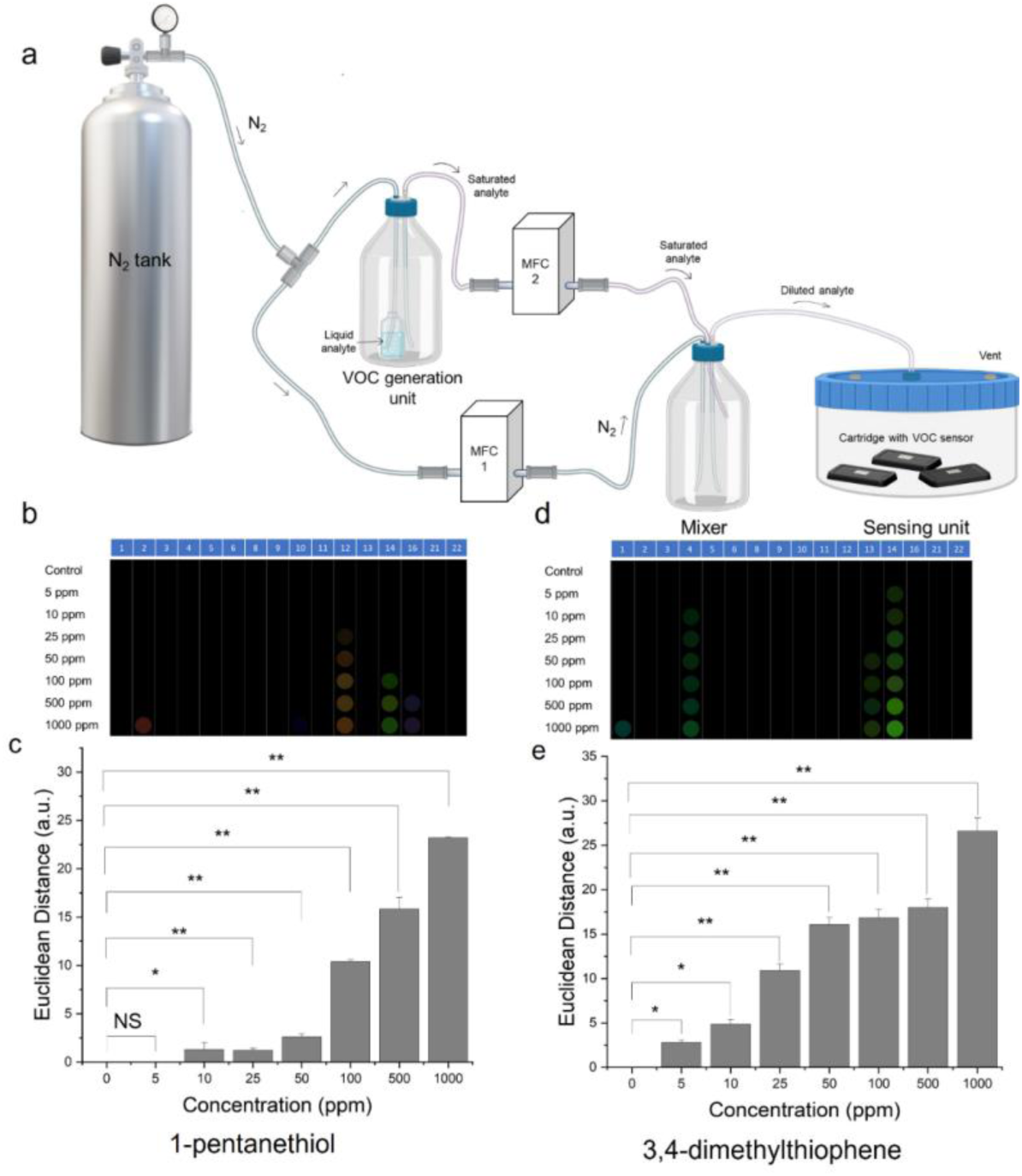
Analytical performance analysis of sensor. (a) Gas mixing rig configuration for the detection of individual volatiles at controlled concentrations; (b) RGB differential colormap of the sensor array at different vapor concentrations of 1-Pentanethiol (n = 3 independent experiments); (c) LOD calculation for detecting 1-Pentathiol vapor (The graph shows statistical significance at p<0.01); (d) RGB differential colormap of the sensor at different vapor concentration of 3,4-dimethylthiophene(n = 3 independent experiments); (e) LOD calculation for detecting 3,4-dimethylthiophene vapor (The graph shows statistical significance at p<0.01(*) and p<0.001(**)). Abbreviations: NS: Not significant, *p<0.01, **p<0.001.

### 3.3 Differentiation of real vegetable samples by the colorimetric sensor array

We tested four different varieties of model vegetables (1A, 2A, 1B, and 2B) from the same species. Three individual sensor measurement experiments were performed for each vegetable sample. The experimental setup is shown in Figure 4a. The sensor responses were collected at 0, 15 min, 30 min, 60 min, and 16h of exposure time. A total of seventeen samples were tested. The left panels of Figure 4b-e show the RGB colormaps of the four varieties of vegetables generated from the images captured by a smartphone-based reader device (**Figure S6**). It was observed that vegetable subclass 1A responded at 15 min of exposure, earlier than other types. However, after 30 min of exposure, significant responses were achieved for subclass 1B and subclass 2A. In the case of subclass 2B, the response time was much longer (∼60 min). Although there was no difference in the physical appearance of the four vegetable varieties, the sensor array successfully differentiated one from another. The progression of color intensity with exposure time is quantitatively demonstrated in right side plots of Figure 4b-e, indicating that 60 min exposure is necessary to observe the strongest response of all vegetable subtypes in this open-air sampling setting. After 16h of incubation, the sensor response patterns become similar to each other due to signal saturation, and therefore not suitable for classification purpose. Figure 4f visually represents the RGB colormaps of four different vegetable varieties tested. The colormap was averaged from three independent measurements for each vegetable sample (total 17 vegetable samples). The unique and distinguishable color pattern observed for each vegetable variety in the colormap indicates that the colorimetric sensor has the ability to differentiate different types of vegetables that belong to the same species. **Figure S3** shows the colormap of all individual samples that were tested.

**Figure 4.**
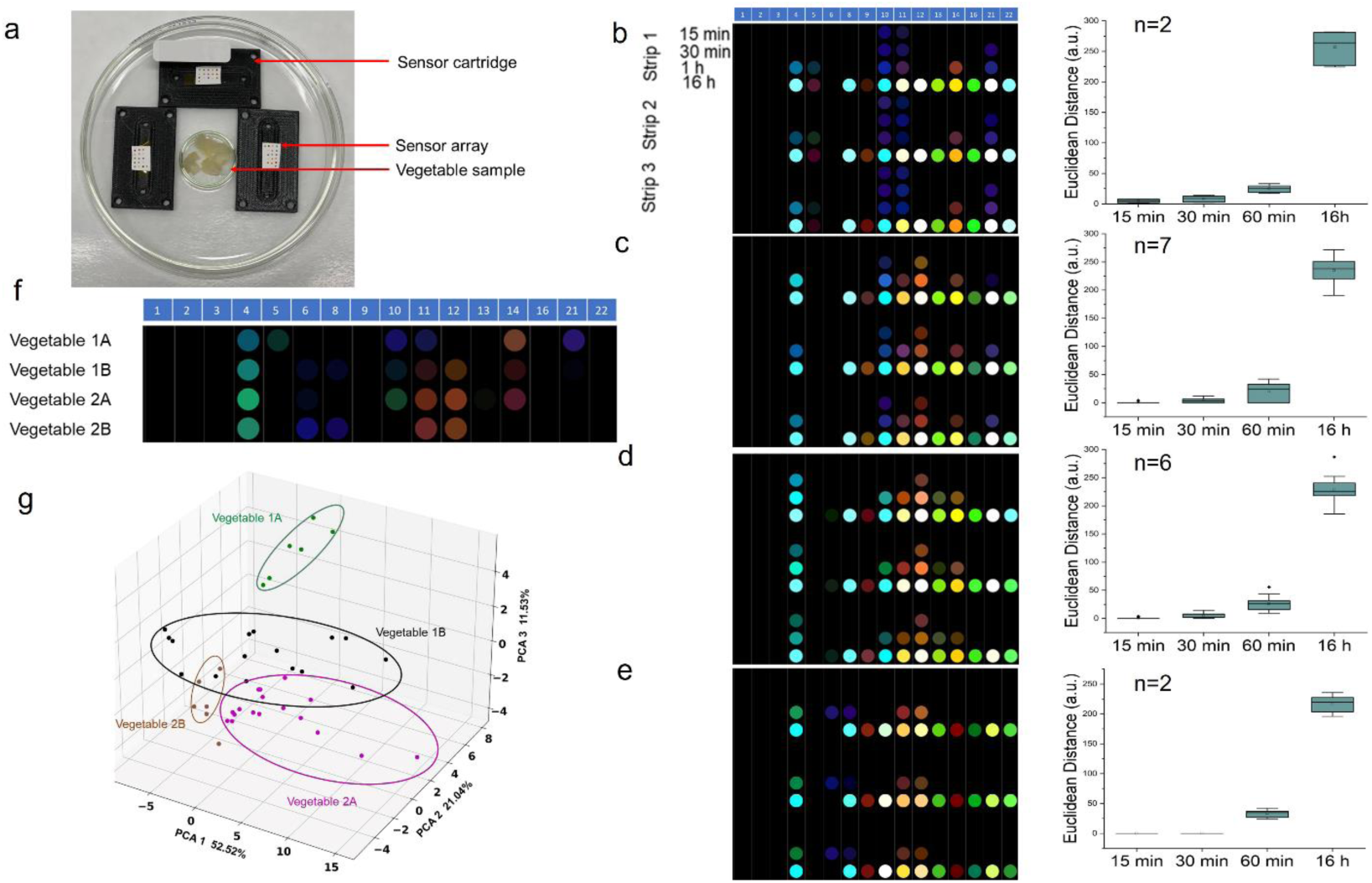
Open-air analysis of real vegetable samples by the colorimetric sensor array. (a) Setup of incubation of sensors with homogenized vegetable samples; (b) RGB profile of vegetable subclass 1A at different exposure time (Strip 1,2,3 show 3 replicates of each sample); (c) RGB profile of vegetable subclass 1B at different exposure time; (d) RGB profile of vegetable subclass 2A at different exposure time; (e) RGB profile of vegetable subclass 2B at different exposure time; (f) Averaged RGB differential colormap of 4 different subtypes of vegetables, showing the differentiable pattern of each vegetable variety (Total: 17 samples, each sample n=3 independent test); (g) PCA plot showing separate clusters for each variety of vegetable.

Moreover, we performed PCA analysis to observe the clustering of sensor response patterns from the different vegetable subclasses. The PCA plot in Figure 4g shows separated clusters for vegetable subclass 1A and subclass 2B. On the other hand, there were a few overlaps between subclass 1B and subclass 2A. We believe a few samples of subclass 1B actually have similar VOC profiles as subclass 2A. Using the first three principal components, we achieved >80% separation.

### 3.4 Needle integration with the colorimetric sensor

For certain vegetables and fruits, the native VOC emission level may be lower than the limit of quantification of the colorimetric sensor array or do not produce certain VOCs until induction. To apply the colorimetric sensor for those zero or low-emitting targets, a dual-functional, needle-integrated device was designed and prototyped. The idea is to use needles to induce VOC emission and subsequent VOC capturing and detection by an adjacent colorimetric sensor array. For a proof-of-concept, we installed syringe needles on a centrifuge tube (50 mL) cap to form the prototype VOC sampling device (Figure 5a). The colorimetric sensor was then attached to the inner middle of the cap. When the needle-loaded cap punched into the vegetable sample, it created tiny holes on the surface of the target sample, which induced VOC emission. The VOCs were then captured by the colorimetric sensor immediately. The cap can also serve as an enclosed chamber for accumulating released VOCs. A smartphone reader was used to capture the image of the sensor strips (Figure 5b). We explored the minimum exposure time needed for a detectable color change on the sensor array (Figure 5c). It was evident that only 2 min exposure time was enough to produce a significantly strong color change of the strips, much shorter than the previous open-air setting (Figure 4a).

**Figure 5.**
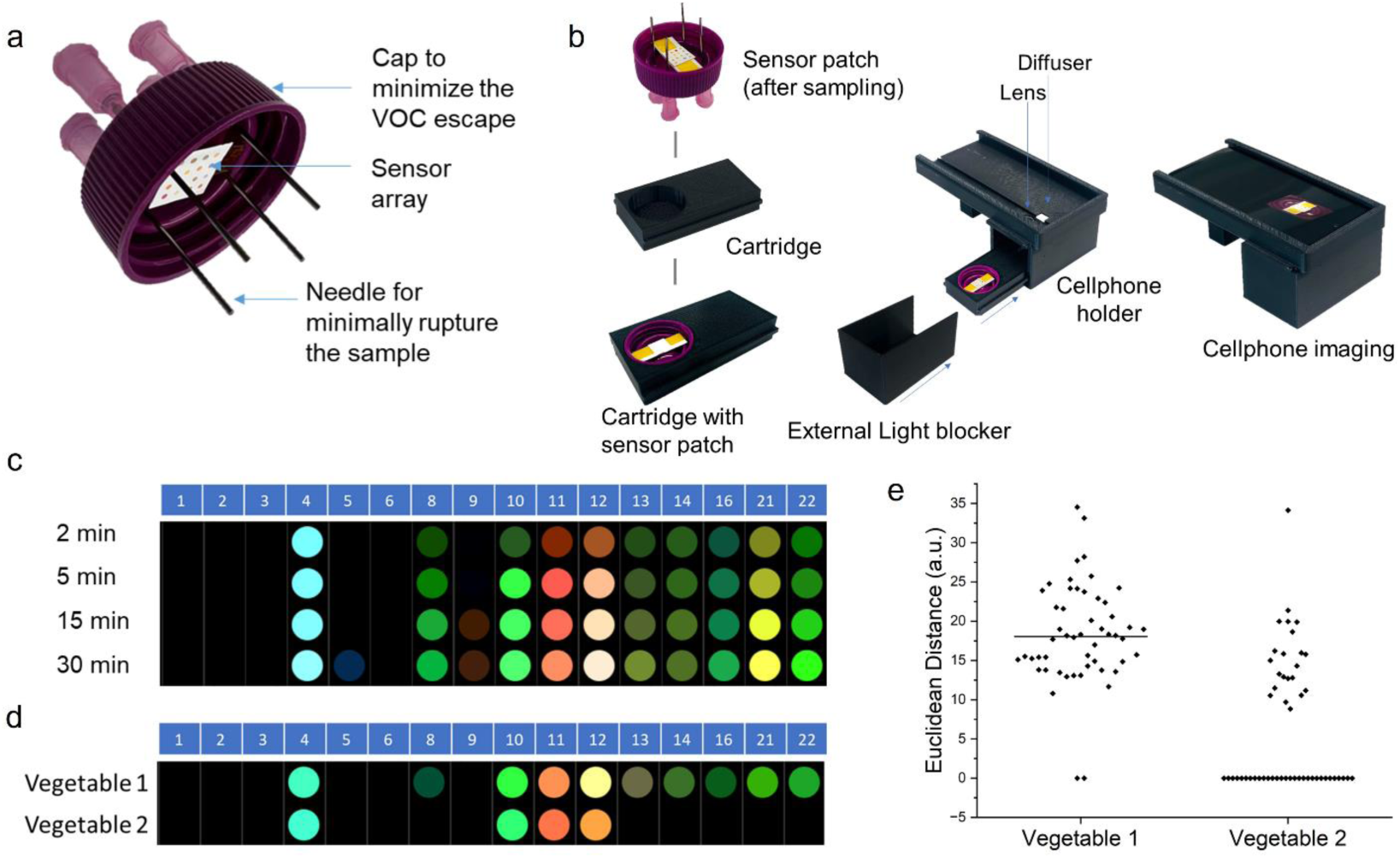
Vegetable variety differentiation using needle-integrated sensor device. (a) A photograph of the actual needle device for punching the surface of vegetables; (b) Schematic of the sensor patch imaging by a smartphone reader; (c) RGB profile of vegetable class 1 punched with needle device with respect to incubation time; (c) RGB profile displaying the difference between vegetable class 1 and class 2; (e) Different sensor response level of class type 1 and type 2.

To discriminate between the vegetable varieties, we conducted an experiment for 2 min exposure of the sensors to a total of 52 vegetable samples, half of which were vegetable class 1 and half class 2. Each vegetable sample was tested twice at two different sampling locations. Figure 5d shows the average differential color patterns between class 1 and class 2. Figure 5e displays the sum of ED values from all 16 sensor spots between two different vegetable varieties, revealing a notable contrast in color change intensity. Sensing dye ID#8, 13, 14, 16, 21, and 22 showed a stronger color difference for the two varieties tested, resulting in a discriminable separation between the two. Interestingly, the sensor responses of class 2 demonstrated a bimodal distribution (Figure 5e). The sensor showed its capability to discern a potentially new subtype of vegetable class 2 that exhibited a similar behavior to class 1. **Figure S5** shows the differential colormap of all 52 vegetable samples with 2 min of exposure.

Along with 50 mL centrifuge tube cap, we also examined several other possible holders to install needles such as water bottle caps, and petri dishes or alternative needle devices such as polymeric microneedle patches. However, all of them produced a strong background color change to the sensors without exposure to vegetable samples (**Figure S4**).

Finally, the performance of the sensor was evaluated by a human-based blind smell test. For the blinded smell test, ten anonymous vegetable samples were tested by personnel who were not involved in sample and sensor preparation. The sample pool contained both vegetable class 1 and 2. The summary of the blind test results is shown in **Table S5**, which confirms the colorimetric sensor results in terms of variety classification.

## 4. Discussion

The newly developed field-portable, minimally invasive VOC detection technology offers significant advantages over conventional gas analysis methods. Unlike GC-MS, which requires expensive equipment, destructive sample preparation methods, and is confined to laboratory settings, our technology is cost-effective, preserves the quality of produce, and allows for on-site analysis with immediate results. Its user-friendly design eliminates the need for specialized training, making it accessible to a broader range of users. Additionally, it has a small footprint and can be easily integrated into various stages of the supply chain, ensuring consistent quality and authenticity checks.

Despite a lot of advantages, there are a few limitations of the current version of the sensing device. For instance, when a new smartphone is employed, re-calibrations are required for the smartphone camera, which may result in varied sensing performance on different phone models. Moreover, extreme environmental conditions (high temperature, humidity, etc.) and the presence of environmental background VOCs can affect the accuracy and consistency of the results, which suggests that the ideal use of VOC strips may be limited in very clean and controlled conditions.(Li et al. 2019) To address the limitations, the future version of optical reader device could be based on a standardizable platform such as Arducam and Raspberry Pi to reduce device-to-device variations and maintain the same level of sensitivity. In addition, designing a robust sensor strip housing to protect it from environmental fluctuations and physical damage will ensure the durability and reliability in various field conditions.

## 5. Conclusion

In summary, we present here a dual-functional needle-integrated VOC sensing platform that is both minimally invasive and field-deployable. This platform has been used to differentiate different vegetable varieties with sufficient detection sensitivity and accuracy in a lab environment. The multiplexed colorimetric sensor array is composed of chemo-responsive and cross-reactive organic dyes encapsulated in hydrophobic ormosil, which allows for the detection and classification of major sulfur-based volatiles, such as thiols, sulfur oxides, and sulfides released from vegetables and fruits. The needles are key components that can minimally invade the vegetable and fruit skins to induce higher concentration VOC emission or generation of characteristic VOCs that do not exist in intact vegetables or fruits. Our research has shown that this platform can successfully differentiate different vegetable varieties in 2 minutes of exposure, which is much quicker than the conventional GC-MS methods. Furthermore, the integrated needle device is cost-effective, with an estimated total cost of <USD 2 (20 cents per colorimetric sensor strip, 5 cents for the cap, and 1.5 USD for the needles). The smartphone attachment (excluding the smartphone) costs ∼USD 15, making this platform an incredibly affordable option compared to currently used detection techniques. This rapid detection can potentially find broad applications in vegetable and fruit quality monitoring and phenotypic classification.

## CRediT authorship contribution statement

**Oindrila Hossain:** Conceptualization, Data Analysis, Investigation, Methodology, Software, Visualization, Writing – original draft, Writing – review & editing; **Yan Wang, Sina Jamalzadegan:** Algorithm development for data processing; **Mingzhuo Li:** Conventional GC-MS Analysis; **Alireza Velayati:** Algorithm development for data processing, Smartphone attachment design; **Noor Mohammad:** Sensitivity analysis, Graphical abstract; **Aditi Dey Poonam:** Microneedle preparation; and **Qingshan Wei:** Conceptualization, Investigation, Writing – review & editing, Funding acquisition, Project administration, Resources, Supervision.

## Acknowledgement

The authors sincerely thank the funding source to conduct the research.

## References

Askim, J.R., Li, Z., LaGasse, M.K., Rankin, J.M., Suslick, K.S., 2016. An optoelectronic nose for identification of explosives. Chemical Science 7(1), 199–206.

Cecchi, L., Ieri, F., Vignolini, P., Mulinacci, N., Romani, A., 2020. Characterization of Volatile and Flavonoid Composition of Different Cuts of Dried Onion (Allium cepa L.) by HS-SPME-GC-MS, HS-SPME-GC×GC-TOF and HPLC-DAD. Molecules.

Cellini, A., Blasioli, S., Biondi, E., Bertaccini, A., Braschi, I., Spinelli, F., 2017. Potential Applications and Limitations of Electronic Nose Devices for Plant Disease Diagnosis. Sensors.

Chatterjee, C., Sen, A., 2015. Sensitive colorimetric sensors for visual detection of carbon dioxide and sulfur dioxide. Journal of Materials Chemistry A 3(10), 5642–5647.

Cuevas, F.J., Moreno-Rojas, J.M., Ruiz-Moreno, M.J., 2017. Assessing a traceability technique in fresh oranges (Citrus sinensis L. Osbeck) with an HS-SPME-GC-MS method. Towards a volatile characterisation of organic oranges. Food chemistry 221, 1930–1938.

Endo, T., Yanagida, Y., Hatsuzawa, T., 2007. Colorimetric detection of volatile organic compounds using a colloidal crystal-based chemical sensor for environmental applications. Sensors and Actuators B: Chemical 125(2), 589–595.

Feng, L., Musto, C.J., Kemling, J.W., Lim, S.H., Zhong, W., Suslick, K.S., 2010. Colorimetric Sensor Array for Determination and Identification of Toxic Industrial Chemicals. Analytical Chemistry 82(22), 9433–9440.

Gao, G., Zhang, X., Yan, Z., Cheng, Y., Li, H., Xu, G., 2022. Monitoring Volatile Organic Compounds in Different Pear Cultivars during Storage Using HS-SPME with GC-MS. Foods 11(23).

Janssens, M., Verlinden, B.E., Eslami Jahromi, K., Pan, Q., Harren, F.J.M., Nicolaï, B.M., Real-time aroma measurements of Conference pears to monitor storage quality using a broadband mid-infrared trace gas sensor. pp. 357–364.

Janzen, M.C., Ponder, J.B., Bailey, D.P., Ingison, C.K., Suslick, K.S., 2006. Colorimetric sensor arrays for volatile organic compounds. Analytical chemistry 78(11), 3591–3600.

Jones, V.P., Horton, D.R., Mills, N.J., Unruh, T.R., Baker, C.C., Melton, T.D., Milickzy, E., Steffan, S.A., Shearer, P.W., Amarasekare, K.G., 2016. Evaluating plant volatiles for monitoring natural enemies in apple, pear and walnut orchards. Biological Control 102, 53–65.

Kangas, M.J., Burks, R.M., Atwater, J., Lukowicz, R.M., Garver, B., Holmes, A.E., 2018. Comparative chemometric analysis for classification of acids and bases via a colorimetric sensor array. Journal of Chemometrics 32(2), e2961.

Lee, G., Hossain, O., Jamalzadegan, S., Liu, Y., Wang, H., Saville, A.C., Shymanovich, T., Paul, R., Rotenberg, D., Whitfield, A.E., Ristaino, J.B., Zhu, Y., Wei, Q., 2023. Abaxial leaf surface-mounted multimodal wearable sensor for continuous plant physiology monitoring. Science Advances 9(15), eade2232.

Leontiev, A.V., Rudkevich, D.M., 2005. Revisiting Noncovalent SO2−Amine Chemistry: An Indicator−Displacement Assay for Colorimetric Detection of SO2. Journal of the American Chemical Society 127(41), 14126–14127.

Li, Z., Liu, Y., Hossain, O., Paul, R., Yao, S., Wu, S., Ristaino, J.B., Zhu, Y., Wei, Q., 2021. Real-time monitoring of plant stresses via chemiresistive profiling of leaf volatiles by a wearable sensor. Matter 4(7), 2553–2570.

Li, Z., Paul, R., Ba Tis, T., Saville, A.C., Hansel, J.C., Yu, T., Ristaino, J.B., Wei, Q., 2019. Non-invasive plant disease diagnostics enabled by smartphone-based fingerprinting of leaf volatiles. Nat Plants 5(8), 856–866.

Lim, S.H., Feng, L., Kemling, J.W., Musto, C.J., Suslick, K.S., 2009. An optoelectronic nose for the detection of toxic gases. Nature Chemistry 1(7), 562–567.

Lim, S.H., Musto, C.J., Park, E., Zhong, W., Suslick, K.S., 2008. A Colorimetric Sensor Array for Detection and Identification of Sugars. Organic Letters 10(20), 4405–4408.

Lindinger, W., Jordan, A., 1998. Proton-transfer-reaction mass spectrometry (PTR–MS): on-line monitoring of volatile organic compounds at pptv levels. Chemical Society Reviews 27(5), 347–375.

Lu, P.-F., Huang, L.-Q., Wang, C.-Z., 2012. Identification and field evaluation of pear fruit volatiles attractive to the oriental fruit moth, Cydia molesta. Journal of chemical ecology 38, 1003–1016.

Mathieu, S., Cin, V.D., Fei, Z., Li, H., Bliss, P., Taylor, M.G., Klee, H.J., Tieman, D.M., 2009. Flavour compounds in tomato fruits: identification of loci and potential pathways affecting volatile composition. Journal of Experimental Botany 60(1), 325–337.

McGorrin, R.J., 2011. The Significance of Volatile Sulfur Compounds in Food Flavors. Volatile Sulfur Compounds in Food, pp. 3-31. American Chemical Society.

Mei, Y., Ge, L., Lai, H., Wang, Y., Zeng, X., Huang, Y., Yang, M., Zhu, Y., Li, H., Li, J., 2023. Decoding the evolution of aromatic volatile compounds and key odorants in Suancai (a Chinese traditional fermented vegetable) during fermentation using stir bar sorptive extraction–gas chromatography–olfactometry–mass spectrometry. LWT 178, 114611.

Monro, T.M., Moore, R.L., Nguyen, M.C., Ebendorff-Heidepriem, H., Skouroumounis, G.K., Elsey, G.M., Taylor, D.K., 2012. Sensing free sulfur dioxide in wine. Sensors (Basel) 12(8), 10759–10773.

Passa, K., Simal, C., Tsormpatsidis, E., Papasotiropoulos, V., Lamari, F.N., 2023. Monitoring of Volatile Organic Compounds in Strawberry Genotypes over the Harvest Period. Plants (Basel) 12(9).

Qiu, S., Wang, J., 2015. Application of sensory evaluation, HS-SPME GC-MS, E-Nose, and E-Tongue for quality detection in citrus fruits. Journal of food science 80(10), S2296–S2304.

Suslick, K.S., Rakow, N.A., Sen, A., 2004. Colorimetric sensor arrays for molecular recognition. Tetrahedron 60(49), 11133–11138.

Taiti, C., Costa, C., Menesatti, P., Caparrotta, S., Bazihizina, N., Azzarello, E., Petrucci, W.A., Masi, E., Giordani, E., 2015. Use of volatile organic compounds and physicochemical parameters for monitoring the post-harvest ripening of imported tropical fruits. European Food Research and Technology 241(1), 91–102.

Taoukis, P., Tsironi, T., 2016. 5 - Smart Packaging for Monitoring and Managing Food and Beverage Shelf Life. In: Subramaniam, P. (Ed.), The Stability and Shelf Life of Food (Second Edition), pp. 141–168. Woodhead Publishing.

Tholl, D., Hossain, O., Weinhold, A., Röse, U.S.R., Wei, Q., 2021. Trends and applications in plant volatile sampling and analysis. The Plant Journal 106(2), 314–325.

Tiwari, S., Kate, A., Mohapatra, D., Tripathi, M.K., Ray, H., Akuli, A., Ghosh, A., Modhera, B., 2020. Volatile organic compounds (VOCs): Biomarkers for quality management of horticultural commodities during storage through e-sensing. Trends in Food Science & Technology 106, 417–433.

Wang, Z., Zhong, T., Chen, X., Yang, B., Du, M., Wang, K., Zalán, Z., Kan, J., 2021. Potential of Volatile Organic Compounds Emitted by Pseudomonas fluorescens ZX as Biological Fumigants to Control Citrus Green Mold Decay at Postharvest. Journal of Agricultural and Food Chemistry 69(7), 2087–2098.

Wu, J., Cao, J., Chen, J., Huang, L., Wang, Y., Sun, C., Sun, C., 2023. Detection and classification of volatile compounds emitted by three fungi-infected citrus fruit using gas chromatography-mass spectrometry. Food Chemistry 412, 135524.

Xiao-wei, H., Zhi-hua, L., Xiao-bo, Z., Ji-yong, S., Han-ping, M., Jie-wen, Z., Li-min, H., Holmes, M., 2016. Detection of meat-borne trimethylamine based on nanoporous colorimetric sensor arrays. Food Chemistry 197, 930–936.

Zhang, H., Xie, Y., Liu, C., Chen, S., Hu, S., Xie, Z., Deng, X., Xu, J., 2017. Comprehensive comparative analysis of volatile compounds in citrus fruits of different species. Food chemistry 230, 316–326.

Zhang, J., Gu, X., Yan, W., Lou, L., Xu, X., Chen, X., 2022. Characterization of Differences in the Composition and Content of Volatile Compounds in Cucumber Fruit. Foods.

Zhang, R., Tang, C., Jiang, B., Mo, X., Wang, Z., 2021a. Optimization of HS-SPME for GC-MS Analysis and Its Application in Characterization of Volatile Compounds in Sweet Potato. Molecules.

Zhang, X., Ren, X., Chingin, K., 2021b. Applications of direct analysis in real time mass spectrometry in food analysis: A review. Rapid Communications in Mass Spectrometry 35(6), e9013.

Zhao, N., Ge, L., Lai, H., Wang, Y., Mei, Y., Huang, Y., Zeng, X., Su, Y., Shi, Q., Li, H., Yuan, H., Zhu, Y., Zuo, Y., Pang, F., Guo, C., Wang, H., Hu, T., 2022. Unraveling the contribution of pre-salting duration to microbial succession and changes of volatile and non-volatile organic compounds in Suancai (a Chinese traditional fermented vegetable) during fermentation. Food Research International 159, 111673.

