## Supporting information for "A Dual-Functional Needle-Based VOC Sensing Platform for Rapid Vegetable Quality Examination"

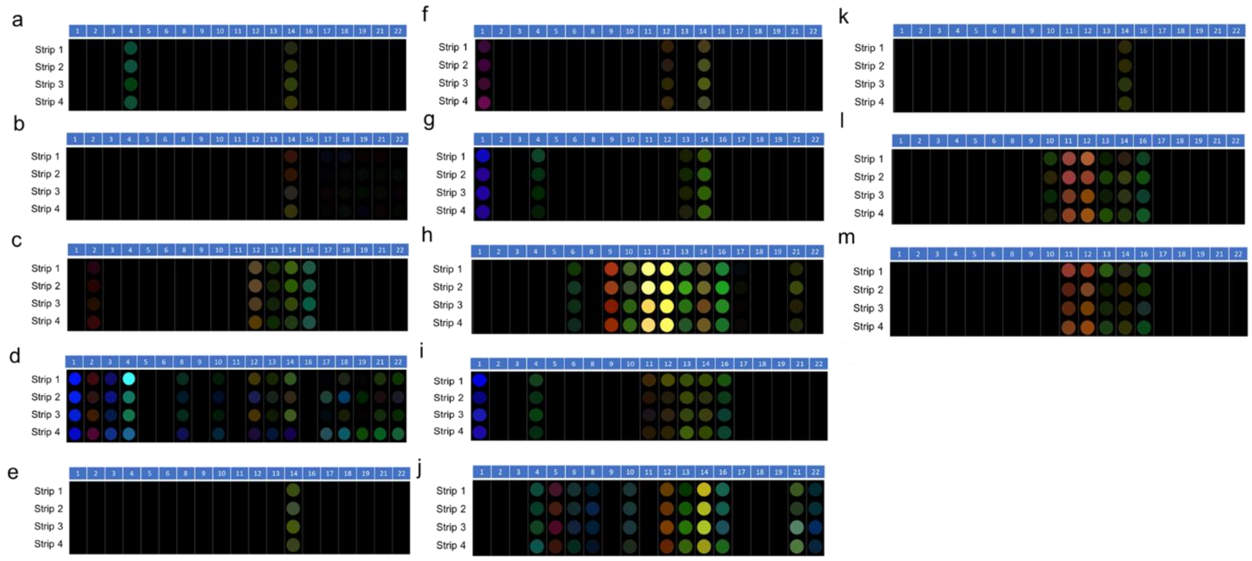


Figure S1. RGB differential colormap of sensors exposed to the vapor of (a) Dimethyl sulfoxide; (b) Propyl sulfoxide; (c) Dimethyl sulfide; (d) Diethyl sulfide; (e) Dipropyl disulfide, (f) Methyl propyl disulfide; (g) 3.4-dimethylthiophene; (h) (E)-2-hexenal; (i) 2-methyl-3-heptanone; (j) Acetone; (k) Methyl salicylate; (l) Methyl jasmonate; (m) 1-octen-3-ol.

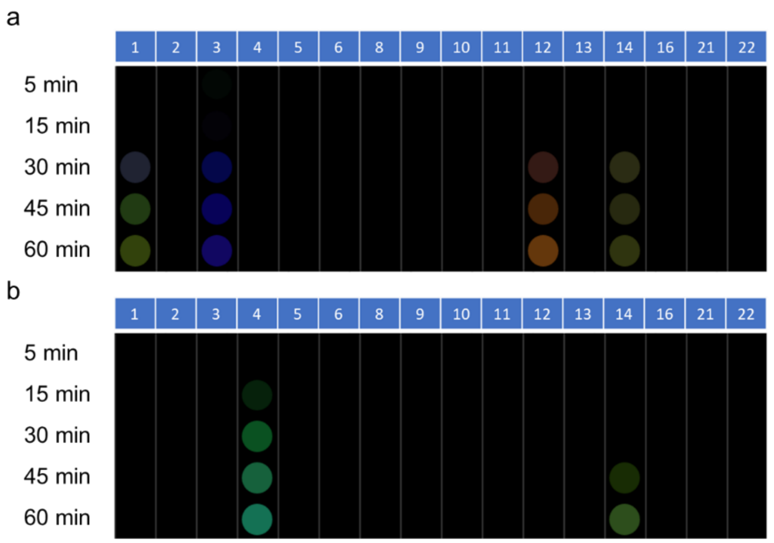


Figure S2. (a) RGB differential profile of sensors exposed to Diethyl sulfide at different exposure time (n=3 independent experiment); (b) RGB profile of sensors exposed to Dimethyl sulfoxide at different exposure time (n=3 independent experiment).

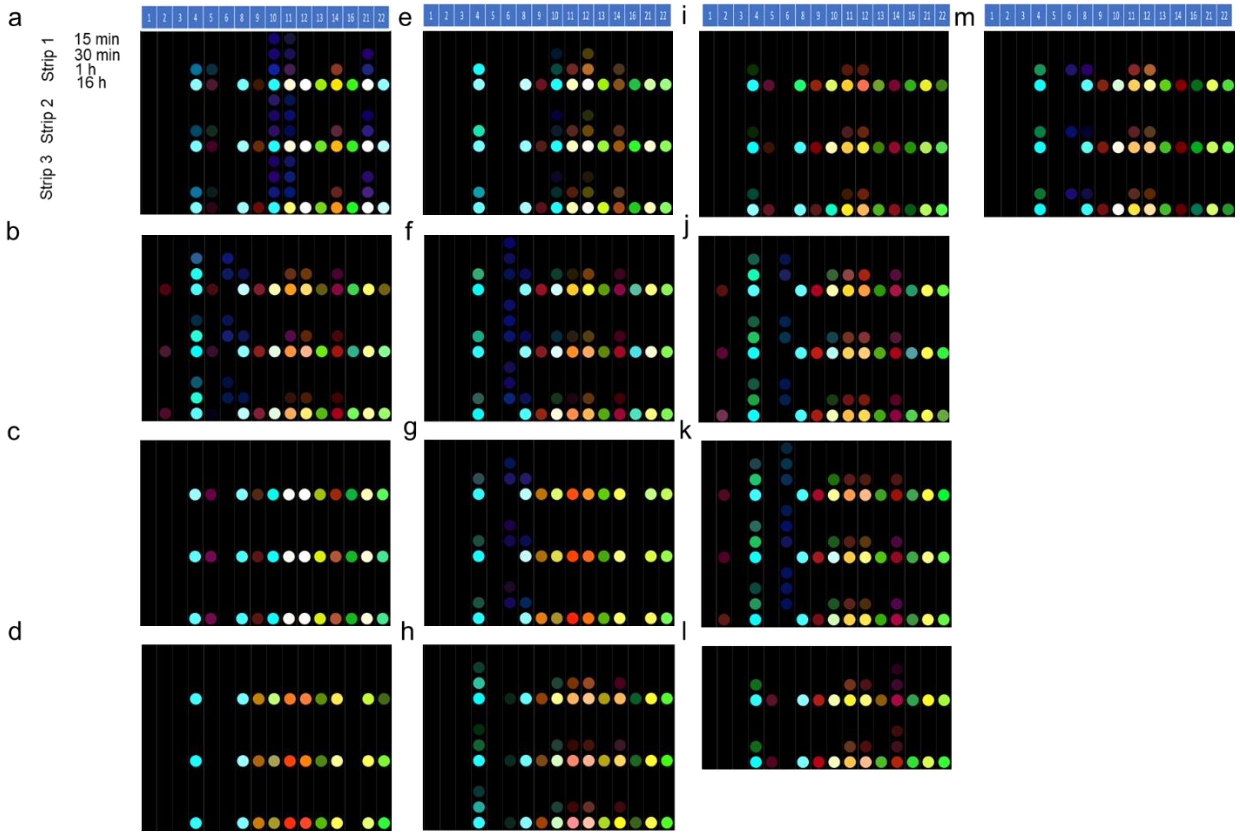


Figure S3. (a) RGB differential profile of vegetable subclass 1A at different exposure time; (b), (c), (d), (e), (f), (g) RGB profile of vegetable subclass 1B at different exposure time, (h), (i), (j), (k), (l) RGB profile of vegetable subclass 2A at different exposure time; (m) RGB profile of vegetable subclass 2B at different exposure time. (Strip 1,2,3 show 3 replicates of each sample).

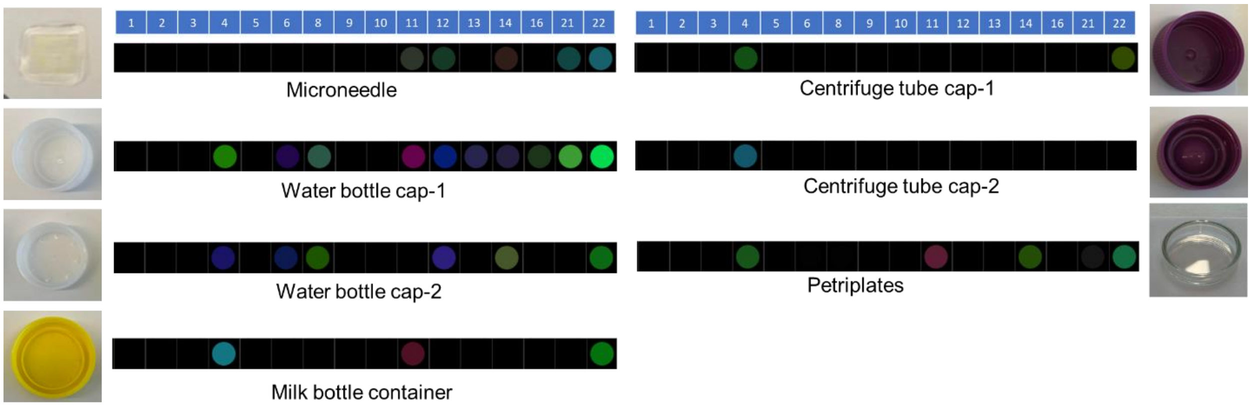


Figure S4. Responses of the different holders to the colorimetric sensor without any VOCs.

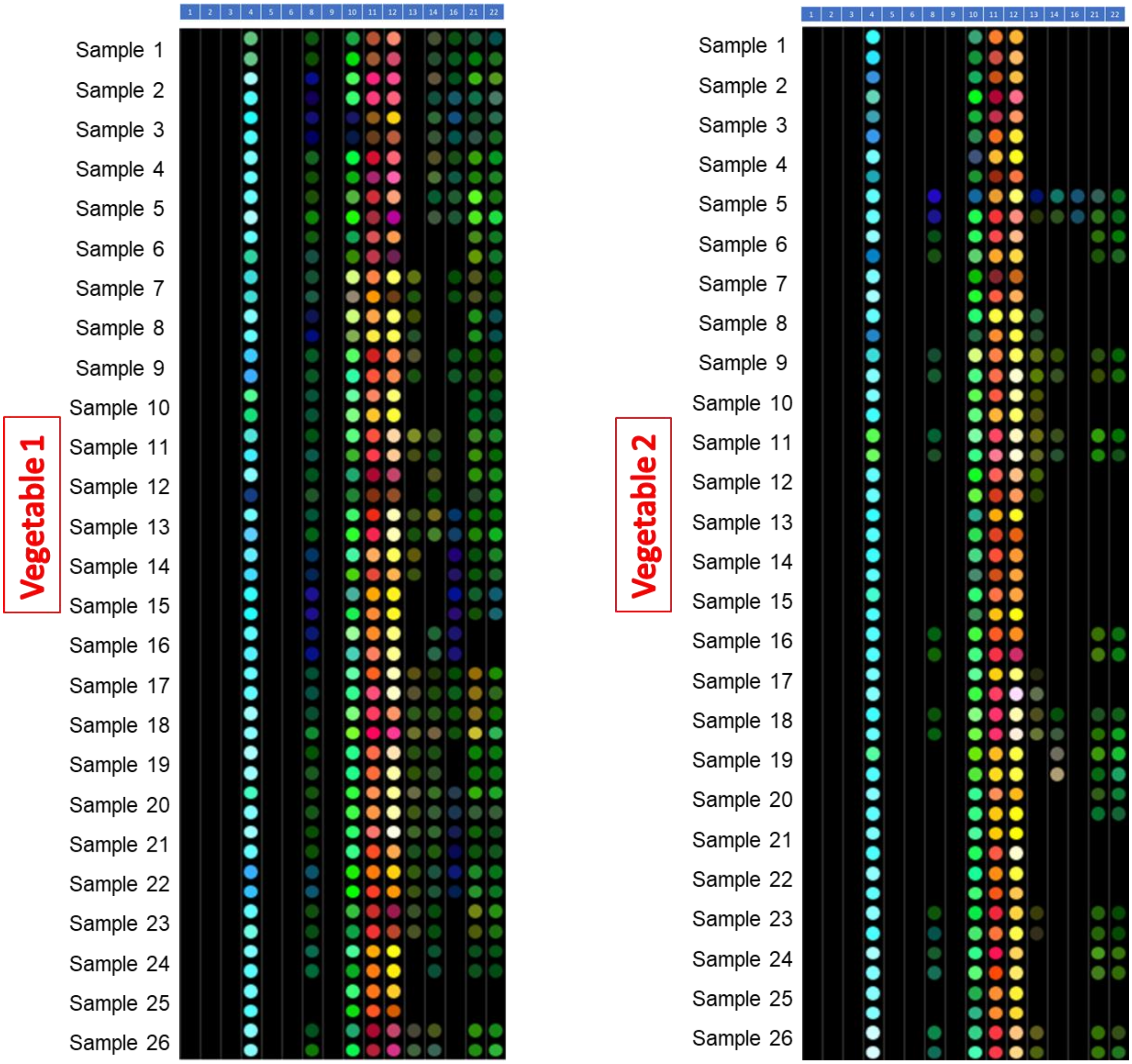


Figure S5. Differential colormap of fifty-two vegetable samples tested with the needle device.

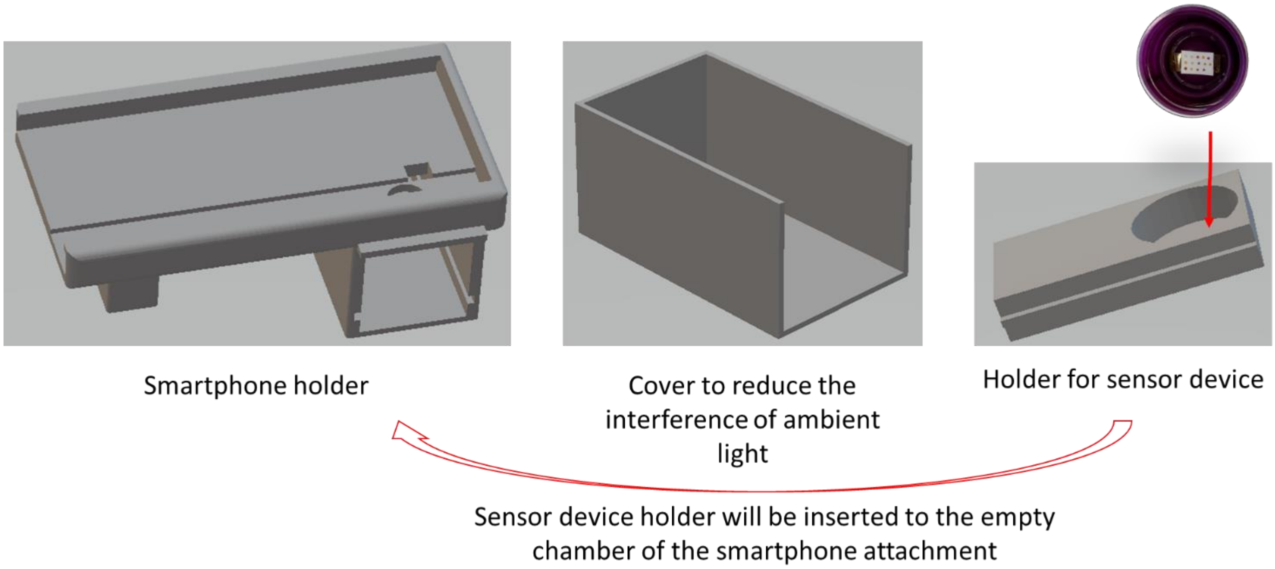


Figure S6. Smartphone-based reader device for capturing images of the colorimetric sensors.

Table S1. Primary selection of colorimetric sensing elements to detect different vegetable-like VOCs.

| **Sensing material components** | **Sensing dye types** |
| --- | --- |
| Zinc-tetraphenylporphyrin Amine<br>Tetrakis(2,4,6-trimethylphenyl)porphyrinatocobalt (II) Amine<br>Tetrakis(pentafluorophenyl)porphine iron(III) chloride Amine | Porphyrins- Amine |
| Methyl Red + sol-gel + TBAH<br>Nitrazine yellow+ sol-gel + TBAH<br>Phenol red + sol-gel + NaOH<br>Cresol red + sol-gel + NaOH | Acid Indicators |
| Pararosaniline hydrochloride + Hydrochloric acid + Formaldehyde + $H_2O$<br>Cresol red + N-methyldiethanolamine (MDEA) | Amine Based |
| $Zn(OAc)_2$ + m-Cresol Purple + sol-gel + TBAH<br>$AgNO_3$ + Bromocresol green + sol-gel + TBAH<br>$AgNO_3$ + Bromophenol Blue + sol-gel + TBAH | Metal Salts |
| Reichardt's dye + sol-gel + TBAH<br>Merocyanine 540 + sol-gel<br>4-(4-nitrobenzyl)pyridine + N- benzyniline + Sol-gel<br>1-ethyl-4-(2-hydroxystyryl)pyridinium iodide + Sol-gel<br>Bromopyrogallol red + Sol-gel | Solvatochromic |
| Thymol blue + sol-gel<br>Bromophenol red + sol-gel<br>Nile red + sol-gel<br>Pyrocatechol Violet + sol-gel<br>Cresol red + sol-gel | Base Indicators |

Table S2. Sensing Dye Response to Vegetable VOC-like Targets from solution-based experiment.

<table>
<tr><th rowspan="2">Sensing material ID no.</th><th rowspan="2">Sensing material components</th><th colspan="6">Target markers</th></tr>
<tr><th>Dimethyl Sulfoxide</th><th>Dipropyl Sulfoxide</th><th>Dimethyl sulfide</th><th>Diethyl sulfide</th><th>(E)-2-Hexenal</th><th>1-Pentanethiol</th></tr>
<tr><td>1</td><td>Zinc-tetraphenylporphyrin Amine</td><td>×</td><td>√</td><td colspan="4" rowspan="9">N/A</td></tr>
<tr><td>2</td><td>Tetrakis(2,4,6-trimethylphenyl)porphyrinatocobalt(II) Amine</td><td>√ (Slight)</td><td>×</td></tr>
<tr><td>3</td><td>Tetrakis(pentafluorophenyl)porphine iron(III) chloride Amine</td><td>×</td><td>√</td></tr>
<tr><td>4</td><td>Methyl Red + sol-gel + NaOH</td><td>×</td><td>√</td></tr>
<tr><td>5</td><td>Nitrazine yellow+ sol-gel + NaOH</td><td>√</td><td>√</td></tr>
<tr><td>6</td><td>Phenol red + sol-gel + NaOH</td><td>×</td><td>√</td></tr>
<tr><td>7</td><td>Cresol red + sol-gel + NaOH</td><td>×</td><td>√</td></tr>
<tr><td>8</td><td>Pararosaniline hydrochloride + Hydrochloric acid + Formaldehyde</td><td>√</td><td>√</td></tr>
<tr><td>9</td><td>Cresol red + Amine</td><td>×</td><td>√</td></tr>
<tr><td>10</td><td>$Zn(OAc)_2$ + m-Cresol Purple + sol-gel + TBAH</td><td>√ (Slight)</td><td>√ (Slight)</td><td colspan="3">N/A</td><td>√</td></tr>
<tr><td>11</td><td>$Zn(OAc)_2$ + Bromocresol green + sol-gel + TBAH</td><td>√ (Slight)</td><td>√ (Slight)</td><td colspan="3">N/A</td><td>√</td></tr>
<tr><td>12</td><td>$Zn(OAc)_2$ + Bromophenol Blue + sol-gel + TBAH</td><td>√ (Slight)</td><td>√ (Slight)</td><td colspan="3">N/A</td><td>√</td></tr>
<tr><td>13</td><td>Reichardt's dye + sol-gel</td><td colspan="2" rowspan="6">N/A</td><td>√</td><td>√</td><td colspan="2" rowspan="6">N/A</td></tr>
<tr><td>14</td><td>Merocyanine 540 + sol-gel</td><td>×</td><td>√</td></tr>
<tr><td>15</td><td>4-(4-nitrobenzyl)pyridine + N-benzyniline + Sol-gel</td><td>√</td><td>√</td></tr>
<tr><td>16</td><td>1-ethyl-4-(2-hydroxystyryl)pyridinium iodide + Sol-gel</td><td>×</td><td>√</td></tr>
<tr><td>17</td><td>Bromopyrogallol red + Sol-gel</td><td>×</td><td>√</td></tr>
<tr><td>18</td><td>Nile red + sol-gel</td><td>√</td><td>√</td></tr>
<tr><td>19</td><td>Thymol blue + sol-gel</td><td>√</td><td colspan="3" rowspan="4">N/A</td><td>√</td><td rowspan="4">N/A</td></tr>
<tr><td>20</td><td>Bromophenol red + sol-gel</td><td>√</td><td>√</td></tr>
<tr><td>21</td><td>Pyrocatechol Violet + sol-gel</td><td>√</td><td>√</td></tr>
<tr><td>22</td><td>Cresol red + sol-gel</td><td>√</td><td>√</td></tr>
</table>

Table S3. Composition of the 22-element colorimetric sensor array.

| Sensing material ID no. | Sensing material components | Amount |
|---|---|---|
| 1 | Zinc-tetraphenylporphyrin + Pyrrolidine (Amine) + Chloroform | 1.36 mg + 1.67 µL + 1 mL |
| 2 | Tetrakis(2,4,6-trimethylphenyl)porphyrinatocobalt (II) + Pyrrolidine (Amine) + Chloroform | 1.58 mg + 1.67 µL + 1 mL |
| 3 | Tetrakis(pentafluorophenyl)porphine iron(III) chloride + Pyrrolidine (Amine) + Chloroform | 2.13 mg + 1.67 µL + 1 mL |
| 4 | Cresol red + Pyrrolidine (Amine) | 0.81 mg + 1.67 µL + 1 mL |
| 5 | Methyl Red + sol-gel + NaOH | 2.5 mg + 500 µL + 5 µL |
| 6 | Nitrazine yellow+ sol-gel + NaOH | 2.5 mg + 500 µL + 0.5 µL |
| 7 | Phenol red + sol-gel + NaOH | 2.5 mg + 500 µL + 0.5 µL |
| 8 | Cresol red + sol-gel + NaOH | 2.5 mg + 500 µL + 1.5 µL |
| 9 | Pararosaniline hydrochloride + Hydrochloric acid + Formaldehyde + $H_2O$ | 1.895 ml (Pararosaniline hydrochloride in 10% ethanol)+ 0.72 ml + 0.0945 ml + 2.2905 mL |
| 10 | $Zn(OAc)_2$ + m-Cresol Purple + sol-gel + TBAH | 3.75 mg + 2.5 mg + 500 µL + 2.5 µL |
| 11 | $Zn(OAc)_2$ + Bromocresol green + sol-gel + TBAH | 3.75 mg + 2.5 mg + 500 µL + 2.5 µL |
| 12 | $Zn(OAc)_2$ + Bromophenol Blue + sol-gel + TBAH | 3.75 mg + 2.5 mg + 500 µL + 2.5 µL |
| 13 | Reichardt's dye + sol-gel | 4.5 mg + 350 µL |
| 14 | Merocyanine 540 + sol-gel | 2 mg + 500 µL |
| 15 | 4-(4-nitrobenzyl)pyridine + N- benzyniline + Sol-gel | 178 mg + 178 mg + 500 µL |
| 16 | 1-ethyl-4-(2-hydroxystyryl)pyridinium iodide + Sol-gel | 3 mg + 500 µL |
| 17 | Bromopyrogallol red + Sol-gel | 2.2 mg + 500 µL |
| 18 | Thymol blue + sol-gel | 2.5 mg + 500 µL |
| 19 | Bromophenol red + sol-gel | 2.5 mg + 500 µL |
| 20 | Nile red + sol-gel | 2.1 mg + 500 µL |
| 21 | Pyrocatechol Violet + sol-gel | 2.2 mg + 500 µL |
| 22 | Cresol red + sol-gel | 2.5 mg + 500 µL |

Red color sensing materials were not used in the final array after the screening.

Table S4. Vapor Pressures of common VOCs at 298 K

| Analyte | Vapor pressure (ppmv) |
|---|---|
| 1-pentanethiol | 18000 |
| Dimethyl sulfoxide | 790 |
| Propyl sulfoxide | Not found |
| Dimethyl sulfide | 660500 |
| Diethyl sulfide | 79000 |
| Dipropyl disulfide | 970 |
| Methyl propyl disulfide | 37763 |
| 3,4-dimethylthiophene | 7690 |
| (E)-2-hexenal | 6080 |
| 2-methyl-3-heptanone | 3325 |
| acetone | 302630 |
| Methyl salicylate | 45 |
| Methyl jasmonate | 0.45 |
| 1-octane-3-ol | 990 |

Vapor pressures data were collected from http://www.thegoodscentscompany.com/index.html, https://www.ncbi.nlm.nih.gov/, or from Material Safety Data Sheets (MSDS); www.msdssearch.com. Due to differences among sources and small variations in temperatures, these values should be taken as approximate only.

Table S5. Blind human test results for classification of vegetable types.

| Vegetables | Sensor test | Blind test | True type |
|---|---|---|---|
| Sample 1 | Vegetable 1 | Vegetable 1 | |
| Sample 2 | Vegetable 1 | Vegetable 1 | |
| Sample 3 | Vegetable 1 | Vegetable 1 | Vegetable 1 |
| Sample 4 | Vegetable 1 | Vegetable 1 | |
| Sample 5 | Vegetable 1 | Vegetable 1 | |
| Sample 6 | Vegetable 2 | Vegetable 2 | |
| Sample 7 | Vegetable 2 | Vegetable 2 | |
| Sample 8 | Vegetable 2 | Vegetable 2 | Vegetable 2 |
| Sample 9 | Vegetable 2 | Vegetable 2 | |
| Sample 10 | Vegetable 2 | Vegetable 2 | |
